## Supplementary file 1 for "The relationship between gut and nasopharyngeal microbiome composition can predict the severity of COVID-19"

|  | **Phyla** | | | |  |  |
| --- | --- | --- | --- | --- | --- | --- |
|  | **Group** | **Actinobacteriota** | **Bacillota** | **Bacteroidota** | **Pseudomonadota** |  |
| **Relative abundance** | **Mild** | 6.938 | 43.003 | 47.362 | 9.403 | **Stool** |
|  | **Moderate** | 4.835 | 34.982 | 59.630 | 7.840 |  |
|  | **Severe** | 8.134 | 39.316 | 45.504 | 11.376 |  |
|  | **Mild** | 60.083 | 30.052 | 34.778 | 41.538 | **Nasopharyngeal** |
|  | **Moderate** | 27.096 | 18.550 | 26.967 | 89.522 |  |
|  | **Severe** | 24.944 | 36.078 | 23.486 | 41.233 |  |
