## Supplementary file 2 for "The relationship between gut and nasopharyngeal microbiome composition can predict the severity of COVID-19"

|  | **Genera** | | | | | | | | | | |
| --- | --- | --- | --- | --- | --- | --- | --- | --- | --- | --- | --- |
|  | **Group** | **Acinetobacter** | **Actinomyces** | **Alcaligenes** | **Alistipes** | **Alloprevotella** | **Anaerococcus** | **Atopobium** | **Bacteroides** | **Campylobacter** | **Corynebacterium** |
| **Relative abundance** | **Mild** | 0.08 | 0.45 | 0.09 | 0.69 | 0.73 | 0.22 | 0.10 | 4.37 | 0.03 | 17.39 |
|  | **Moderate** | 0.08 | 0.02 | 0.19 | 0.15 | 0.13 | 0.01 | 0.01 | 3.45 | 0.00 | 0.20 |
|  | **Severe** | 0.14 | 0.90 | 0.05 | 0.04 | 0.97 | 0.39 | 0.22 | 0.24 | 0.09 | 10.94 |
|  | **a p-value** | 0.04 | 0.0004 | - | 0.0258 | - | 0.0027 | 0.0014 | 0.0072 | 0.0004 | 0.0016 |
|  | **b p-value** | 0.018 | 0.0001 | 0.0004 | - | 0.0004 | 0.0001 | 0.0001 | 0.0001 | 0.0001 | 0.0001 |
|  | **c p-value** | - | - | 0.0001 | 0.032 | 0.0001 | 0.0091 | - | 0.005 | - | - |
|  | **Genera** | | | | | | | | | | |
|  | **Group** | **Cutibacterium** | **Dolosigranulum** | **Enterobacter** | **Enterococcus** | **Finegoldia** | **Fusobacterium** | **Gemella** | **Haemophilus** | **Lachnoanaerobaculum** | **Lachnospiraceae_NK4A136_group** |
| **Relative abundance** | **Mild** | 3.51 | 0.25 | 0.23 | 1.07 | 0.07 | 0.23 | 0.12 | 0.20 | 0.06 | 3.86 |
|  | **Moderate** | 0.23 | 0.00 | 0.22 | 0.03 | 0.01 | 0.02 | 0.01 | 0.06 | 0.00 | 0.13 |
|  | **Severe** | 4.25 | 0.56 | 2.08 | 6.56 | 0.10 | 1.27 | 0.25 | 0.79 | 0.41 | 0.02 |
|  | **a p-value** | 0.009 | - | 0.0005 | 0.0029 | 0.0041 | 0.0167 | 0.0098 | 0.0045 | 0.0004 | - |
|  | **b p-value** | 0.0001 | 0.0231 | 0.0001 | 0.0001 | 0.0001 | 0.0001 | 0.0001 | 0.0001 | - | 0.0008 |
|  | **c p-value** | 0.0052 | - | - | 0.0282 | - | 0.0487 | - | 0.0041 | 0.0017 | 0.0001 |
|  | **Genera** | | | | | | | | | | |
|  | **Group** | **Lactobacillus** | **Lawsonella** | **Leptotrichia** | **Megasphaera** | **Muribaculaceae** | **Neisseria** | **Oribacterium** | **P5D1-392** | **Peptoniphilus** | **Porphyromonas** |
| **Relative abundance** | **Mild** | 4.61 | 0.56 | 0.08 | 0.03 | 14.10 | 0.35 | 0.10 | 0.16 | 0.24 | 0.66 |
|  | **Moderate** | 0.84 | 0.02 | 0.01 | 0.01 | 0.69 | 0.04 | - | - | - | - |
|  | **Severe** | 1.13 | 0.22 | 1.01 | 0.16 | 0.21 | 0.62 | - | - | - | - |
|  | **a p-value** | 0.0285 | 0.0012 | 0.0021 | 0.0148 | 0.0001 | - | - | - | - | - |
|  | **b p-value** | 0.0001 | 0.0001 | 0.0001 | - | 0.047 | - | - | - | - | - |
|  | **c p-value** | - | - | - | 0.0425 | 0.0001 | 0.0121 | - | - | - | - |
|  | **Group** | **Prevotella** | **Pseudarthrobacter** | **Pseudomonas** | **Rothia** | **Serratia** | **Staphylococcus** | **Stenotrophomonas** | **Streptococcus** | **Un. Comamonadaceae** | **Un. Lachnospiraceae** |
| **Relative abundance** | **Mild** | 1.57 | - | - | - | - | - | - | 1.2 | - | 0.68 |
|  | **Moderate** | 1.29 | 1.93 | 74.4 | - | - | 0.11 | 0.26 | 0.34 | 0.37 | - |
|  | **Severe** | 9.06 | - | 17.03 | 0.72 | 3.92 | 9.9 | 3.66 | 2.83 | 0.57 | - |
|  | **a p-value** | 0.0002 | - | - | - | - | - | - | 0.0005 | - | - |
|  | **b p-value** | - | - | - | - | - | - | - | - | - | - |
|  | **c p-value** | 0.0001 | - | 0.00029 | - | - | 0.00017 | 0.000538 | 0.0001 | 0.09 | - |
|  | **Group** | **Veillonella** |  | | | | | | | | |
| **Relative abundance** | **Mild** | - |  |  |  |  |  |  |  |  |  |
|  | **Moderate** | - |  |  |  |  |  |  |  |  |  |
|  | **Severe** | 4.77 |  |  |  |  |  |  |  |  |  |
|  | **a p-value** | - |  |  |  |  |  |  |  |  |  |
|  | **b p-value** | - |  |  |  |  |  |  |  |  |  |
|  | **c p-value** | - |  |  |  |  |  |  |  |  |  |
