## Supplementary file 3 for "The relationship between gut and nasopharyngeal microbiome composition can predict the severity of COVID-19"

|  | **Genera** | | | | | | | | | | |
| --- | --- | --- | --- | --- | --- | --- | --- | --- | --- | --- | --- |
|  | **Group** | **[Eubacterium]_coprostanoligenes_group** | **Alistipes** | **Anaerococcus** | **Bacteroides** | **Barnesiella** | **Blautia** | **Christensenellaceae_R-7_group** | **Clostridia_UCG-014** | **Coprococcus** | **Dialister** |
| **Relative abundance** | **Mild** | 1.50 | 1.83 | 0.04 | 30.04 | 0.90 | 2.60 | 1.08 | 1.64 | 0.68 | 0.24 |
|  | **Moderate** | 1.14 | 2.42 | 0.02 | 38.12 | 1.05 | 1.87 | 1.84 | 0.65 | 0.36 | 0.14 |
|  | **Severe** | 0.71 | 5.39 | 1.55 | 16.64 | 0.24 | 0.84 | 0.52 | 0.15 | 0.17 | 1.97 |
|  | **a p-value** | - | - | - | - | - | - | - | - | - | - |
|  | **b p-value** | 0.000148774 | - | 9.14443E-11 | 0.010104947 | 0.0037286 | - | - | 4.50596E-07 | 1.5262E-05 | 1.62114E-05 |
|  | **c p-value** | 0.000465315 | - | 4.7502E-15 | 2.46458E-06 | 0.004409355 | 0.000175237 | 0.00416827 | 7.73712E-06 | 0.001837231 | 1.49617E-08 |
|  | **Genera** | | | | | | | | | | |
|  | **Group** | **Dorea** | **Faecalibacterium** | **Finegoldia** | **Lachnoclostridium** | **Lachnospiraceae_NK4A136_group** | **Muribaculaceae** | **Parabacteroides** | **Peptoniphilus** | **Prevotella** | **Roseburia** |
| **Relative abundance** | **Mild** | 0.58 | 5.39 | 0.07 | 0.44 | 0.76 | 2.66 | 2.92 | 0.04 | 4.04 | 0.65 |
|  | **Moderate** | 0.56 | 5.38 | 0.04 | 0.70 | 0.43 | 0.47 | 5.05 | 0.03 | 9.75 | 0.65 |
|  | **Severe** | 0.15 | 0.77 | 2.36 | 2.74 | 0.75 | 0.50 | 2.81 | 1.24 | 17.53 | 0.37 |
|  | **a p-value** | - | - | - | - | 0.002921415 | - | 0.024609908 | - | - | - |
|  | **b p-value** | 5.43251E-06 | 1.56601E-06 | 4.01545E-11 | 1.06877E-07 | 0.046073962 | 2.39299E-06 | - | 4.01419E-11 | 8.29752E-05 | 0.015758969 |
|  | **c p-value** | 3.07997E-05 | 2.79925E-09 | 1.23129E-14 | 8.92379E-07 | - | 0.000775785 | 0.003882489 | 6.07343E-15 | 0.001165821 | 0.049197388 |
|  | **Genera** | | | | | | | | | | |
|  | **Group** | **Ruminococcus** | **Streptococcus** | **Subdoligranulum** | **UCG-002** | **Unclassified Lachnospiraceae** |  | | | | |
| **Relative abundance** | **Mild** | 1.04 | 0.59 | 1.16 | 1.42 | 3.54 |  | | | | |
|  | **Moderate** | 0.48 | 2.04 | 0.73 | 1.67 | 1.99 |  |  |  |  |  |
|  | **Severe** | 0.13 | 1.30 | 0.32 | 1.70 | 1.02 |  |  |  |  |  |
|  | **a p-value** | - | 0.003059416 | - | - | 0.031207777 |  |  |  |  |  |
|  | **b p-value** | 3.67848E-06 | 0.001860069 | 0.000191191 | - | 1.20879E-05 |  |  |  |  |  |
|  | **c p-value** | 2.34358E-05 | - | 0.001599171 | - | 0.007282069 |  |  |  |  |  |
