## Supplementary file 4 for "The relationship between gut and nasopharyngeal microbiome composition can predict the severity of COVID-19"

| **Nasopharyngeal** | | | **Stool** | | |
| --- | --- | --- | --- | --- | --- |
| **Mild** | **Moderate** | **Severe** | **Mild** | **Moderate** | **Severe** |
| *Haemophilus sp.* | *Achromobacter spanius* | *Paenarthrobacter nitroguajacolicus* | *Dialister pneumosintes* | *Anaerotruncus sp.* | *Bacteroides plebeius* |
| *Lactobacillus fermentum* | *Bacillus coagulans* | *Bacteroides uniformis* | *Anaerococcus prevotii* | *Peptoniphilus gorbachii* | *Coprobacillus cateniformis* |
| *Lactobacillus reuteri* | *Prevotella nanceiensis* | *Xanthomonas sp.* | *Fusobacterium sp.* | *Peptoniphilus duerdenii* | *Alistipes indistinctus* |
| *Prevotella nigrescens* | *Corynebacterium freneyi* | *Citrobacter europaeus* | *Bacteroides ovatus* | *Varibaculum cambriense* | *Alistipes inops* |
| *Peptostreptococcus stomatis* | *Acinetobacter johnsonii* |  | *Peptostreptococcus stomatis* | *Bacteroides eggerthii* | *Anaerotignum lactatifermentans* |
| *Burkholderia sp.* | *Bacteroides vulgatus* |  | *Bacteroides coagulans* |  | *Anaerotruncus colihominis* |
| *Actinomyces graevenitzii* | *Dialister propionicifaciens* |  | *Porphyromonas bennonis* |  | *Blautia hansenii* |
| *Prevotella pallens* | *Burkholderia sp.* |  | *Anaerococcus octavius* |  | *Bifidobacterium bifidum* |
| *Prevotella sahii* | *Lactobacillus gasseri* |  |  |  | *Peptoniphilus urinimassiliensis* |
| *Capnocytophaga granulosa* | *Prevotella melaninogenica* |  |  |  | *Prevotella stercorea* |
| *Prevotella histicola* |  |  |  |  |  |
| *Alloprevotella tannerae* |  |  |  |  |  |
| *Prevotella oris* |  | | | | |
