## Supplementary file 5 for "The relationship between gut and nasopharyngeal microbiome composition can predict the severity of COVID-19"

| **Group** | **Species** | **Relative Abundance** |
| --- | --- | --- |
| Mild | *Alistipes onderdonkii* | 0.23 |
| Mild | *Bacteroides cellulosilyticus* | 0.31 |
| Mild | *Bacteroides coprocola* | 1.75 |
| Mild | *Bacteroides stercoris* | 1.02 |
| Mild | *Escherichia sp.* | 2.18 |
| Mild | *Prevotella bivia* | 0.04 |
| Mild | *Prevotella copri* | 3.06 |
| Mild | *Prevotella stercorea* | 0.07 |
| Mild | *Prevotella timonensis* | 0.01 |
| Mild | *Ruminococcus bicirculans* | 0.51 |
| Mild | *Streptococcus salivarius* | 0.49 |
| Mild | *Sutterella stercoricanis* | 0.28 |
| Mild | *Veillonella ratti* | 0.54 |
| Moderate | *Alistipes onderdonkii* | 0.48 |
| Moderate | *Bacteroides cellulosilyticus* | 1.76 |
| Moderate | *Bacteroides coprocola* | 1.73 |
| Moderate | *Bacteroides stercoris* | 2.63 |
| Moderate | *Escherichia sp.* | 0.39 |
| Moderate | *Prevotella bivia* | 0.03 |
| Moderate | *Prevotella copri* | 5.99 |
| Moderate | *Prevotella stercorea* | 1.24 |
| Moderate | *Prevotella timonensis* | 0.04 |
| Moderate | *Ruminococcus bicirculans* | 0.30 |
| Moderate | *Streptococcus salivarius* | 1.78 |
| Moderate | *Sutterella stercoricanis* | 0.09 |
| Moderate | *Veillonella ratti* | 0.01 |
| Severe | *Alistipes onderdonkii* | 3.10 |
| Severe | *Bacteroides cellulosilyticus* | 0.42 |
| Severe | *Bacteroides coprocola* | 0.11 |
| Severe | *Bacteroides stercoris* | 0.33 |
| Severe | *Escherichia sp.* | 2.75 |
| Severe | *Prevotella bivia* | 7.12 |
| Severe | *Prevotella copri* | 0.28 |
| Severe | *Prevotella stercorea* | 0.03 |
| Severe | *Prevotella timonensis* | 4.38 |
| Severe | *Ruminococcus bicirculans* | 0.05 |
| Severe | *Streptococcus salivarius* | 0.08 |
| Severe | *Sutterella stercoricanis* | 0.02 |
| Severe | *Veillonella ratti* | 0.02 |
