## Supplementary file 6 for "The relationship between gut and nasopharyngeal microbiome composition can predict the severity of COVID-19"

| **Group** | **Species** | **Relative Abundance** |
| --- | --- | --- |
| Mild | *Azotobacter chroococcum* | 0.012 |
| Mild | *Burkholderia glumae* | 0.237 |
| Mild | *Haemophilus parainfluenzae* | 0.162 |
| Mild | *Leptotrichia sp.* | 0.015 |
| Mild | *Massilia niabensis* | 0.013 |
| Mild | *Metamycoplasma salivarium* | 0.006 |
| Mild | *Paraburkholderia caledonica* | 0.015 |
| Mild | *Prevotella dentalis* | 0.014 |
| Mild | *Pseudomonas veronii* | 0.006 |
| Mild | *Stenotrophomonas rhizophila* | 0.010 |
| Moderate | *Azotobacter chroococcum* | 0.457 |
| Moderate | *Burkholderia glumae* | 0.176 |
| Moderate | *Haemophilus parainfluenzae* | 0.052 |
| Moderate | *Leptotrichia sp.* | 0.000 |
| Moderate | *Massilia niabensis* | 0.001 |
| Moderate | *Metamycoplasma salivarium* | 0.002 |
| Moderate | *Paraburkholderia caledonica* | 0.000 |
| Moderate | *Prevotella dentalis* | 0.000 |
| Moderate | *Pseudomonas veronii* | 0.164 |
| Moderate | *Stenotrophomonas rhizophila* | 0.207 |
| Severe | *Azotobacter chroococcum* | 0.022 |
| Severe | *Burkholderia glumae* | 0.105 |
| Severe | *Haemophilus parainfluenzae* | 0.641 |
| Severe | *Leptotrichia sp.* | 0.213 |
| Severe | *Massilia niabensis* | 0.011 |
| Severe | *Metamycoplasma salivarium* | 0.016 |
| Severe | *Paraburkholderia caledonica* | 0.008 |
| Severe | *Prevotella dentalis* | 0.092 |
| Severe | *Pseudomonas veronii* | 0.089 |
| Severe | *Stenotrophomonas rhizophila* | 0.002 |
