## Supplementary file 7 for "The relationship between gut and nasopharyngeal microbiome composition can predict the severity of COVID-19"

| **Species** | **Clinical variable** | **Correlation value** | **P.value** | **Sample** |
| --- | --- | --- | --- | --- |
| *Prevotella timonensis* | Age | 0.433220642 | 0.0001 | Stool |
| *Alistipes onderdonkii* | Dyspnoea | -0.302873618 | 4.8082E-237 | Stool |
| *Prevotella timonensis* | Dyspnoea | -0.261490563 | 4.7871E-175 | Stool |
| *Prevotella copri* | RF | -0.316747263 | 2.913E-260 | Stool |
| *Bacteroides coprocola* | RF | -0.30959655 | 3.8748E-248 | Stool |
| *Prevotella timonensis* | RF | 0.401670518 | 0.0001 | Stool |
| *Ruminococcus bicirculans* | RF | -0.390957133 | 0.0001 | Stool |
| *Sutterella stercoricanis* | RF | -0.318312438 | 5.8285E-263 | Stool |
| *Prevotella bivia* | Cardiomyopathy | -0.349922087 | 0.0001 | Stool |
| *Escherichia sp.* | Cardiomyopathy | -0.25645094 | 3.2642E-168 | Stool |
| *Alistipes onderdonkii* | Cardiomyopathy | -0.266597375 | 3.9795E-182 | Stool |
| *Prevotella timonensis* | Cardiomyopathy | -0.33913253 | 1.9787E-300 | Stool |
| *Veillonella sp.* | sPO2 | -0.262648324 | 1.2265E-176 | Stool |
| *Prevotella bivia* | Lymphocytes | -0.452509034 | 0.0001 | Stool |
| *Escherichia sp.* | Lymphocytes | -0.301124524 | 3.3133E-234 | Stool |
| *Prevotella timonensis* | Lymphocytes | -0.400188705 | 0.0001 | Stool |
| *Prevotella bivia* | CRP | 0.499050333 | 0.0001 | Stool |
| *Prevotella copri* | CRP | -0.354463929 | 0.0001 | Stool |
| *Bacteroides coprocola* | CRP | -0.264543179 | 2.929E-179 | Stool |
| *Prevotella timonensis* | CRP | 0.543676913 | 0.0001 | Stool |
| *Veillonella sp.* | CRP | 0.421802104 | 0.0001 | Stool |
| *Ruminococcus bicirculans* | CRP | -0.374671876 | 0.0001 | Stool |
| *Sutterella stercoricanis* | CRP | -0.346312313 | 0.0001 | Stool |
| *Burkholderia glumae* | Lymphocytes | -0.263526324 | 7.5241E-178 | Nasal swab |
| *Leptotrichia sp.* | Lymphocytes | -0.25403711 | 5.4113E-165 | Nasal swab |
| *Prevotella dentalis* | Lymphocytes | -0.259633949 | 1.6417E-172 | Nasal swab |
| *Metamycoplasma salivarium* | Lymphocytes | -0.260736158 | 5.1607E-174 | Nasal swab |
